# Social attraction mediates collective foraging decisions in invasive hornets

**DOI:** 10.1101/2025.11.20.689504

**Authors:** Mathilde Lacombrade, Charlotte Doussot, Blandine Mahot-Castaing, Jacques Gautrais, Loïc Goulefert, Edgar Ferrieu, Amelie Bourg, Denis Thiéry, Fanny Vogelweith, Mathieu Lihoreau

## Abstract

Group-living animals commonly use social information to better locate and exploit resources. In many insects, birds, fish and mammals, this can lead to collective foraging decisions by which animals share a single food source among alternatives of equal qualities. Here, we report collective foraging decisions in a social wasp, the yellow-legged hornet *Vespa velutina nigrithorax*, a major predator of bees and other terrestrial invertebrates invasive across Asia, Europe and North America. When given a choice between two identical liquid food sources (feeders or traps containing sugar solutions), wild hornets distributed asymmetrically on the two options, and this phenomenon was more frequent as group size increased. Priming one of the food sources with dead hornets predictably biased the collective choices towards this particular option, irrespective of whether the dead insects were conspecifics or hornets from a closely related species. Inter-attraction in yellow-legged hornets is thus a passive and non-specific mechanism, possibly mediated by visual or chemical cues displayed by dead hornets. This collective behaviour may provide important foraging advantages to hornets invading new territories and bring new perspectives for population control.

## Introduction

Social animals often make collective decisions when it comes to fleeing danger, choosing a new home, or finding food [1]. These collective behaviours can involve dozens to billions of individuals [2] and provide fitness benefits through swarm intelligence [3–5]. Strikingly, this rich diversity of collective phenomena often emerge from a limited number of generic principles, such as feedback loops and quorum sensing by which the behaviour of some individuals in the group influences that of others in a non-linear manner [6]. For instance, when foraging for food, many ant species deposit pheromone trails that attract nestmates on site. The newcomers then amplify the signal, ultimately recruiting more ants. When a sufficient number of foragers are engaged in the task, pheromone deposition acts as a positive feedback loop that leads the colony to exploit one food source in particular among several alternatives [7]. If the food sources vary in quality, the ants can collectively select the most profitable one [8]. Conversely, food depletion and pheromone degradation constitute negative feedbacks that can reduce or stop the expression of the collective behaviour [8].

Collective decisions can be experimentally demonstrated by giving an animal group a choice between two identical options, as for instance two food sites of equivalent size and nutrient content [8,9]. If the animals use social cues to decide where to go, feedback loops should result in an asymmetrical distribution of individuals on the two options [7]. The strength of the collective decision, as measured by the degree of asymmetry, typically increases with group size (i.e. the number of individuals involved in foraging) and the amount of social cues available [10]. Importantly, while several animals make collective decisions through active signaling (e.g. ant pheromone trail to food sources [11], honeybees waggle dance advertising feeding location [12]), asymmetrical distributions can also emerge through passively displayed social cues (e.g. body scents of cockroaches choosing a resting site [13]).

Here, we explored the possibility for collective decision-making in an ecologically and economically important social wasp, the yellow-legged hornet (*Vespa velutina nigrithorax*). In recent years, this hornet native of South Eastern Asia, has become invasive across Western Europe [14], Eastern Asia [15,16] and Northern America [17]. In these areas, hornets feed from flower nectar as a source of energy [18,19]. However, they also cause strong concerns for local invertebrate biodiversity and beekeeping activities. Honeybees, in particular, account for 33 to 66% of their diet [20,21], leading many beekeepers to use handcrafted traps baited with sugary food to control populations [22]. Using this method to protect our own experimental apiaries, we observed that similar traps placed on the same site often caught very different numbers of hornets, suggesting the occurrence of social attraction between hornets. To test this hypothesis, we gave wild hornets a choice between two identically attractive food sources placed next to each other (two feeders containing diluted honey or two traps containing wine, beer and sugar; Figure 1) and quantified their distribution through time. If the hornets attracted each other, we expected them to be distributed asymmetrically between the two options, and the frequency of significant asymmetries to increase with group size.

**Figure 1:**
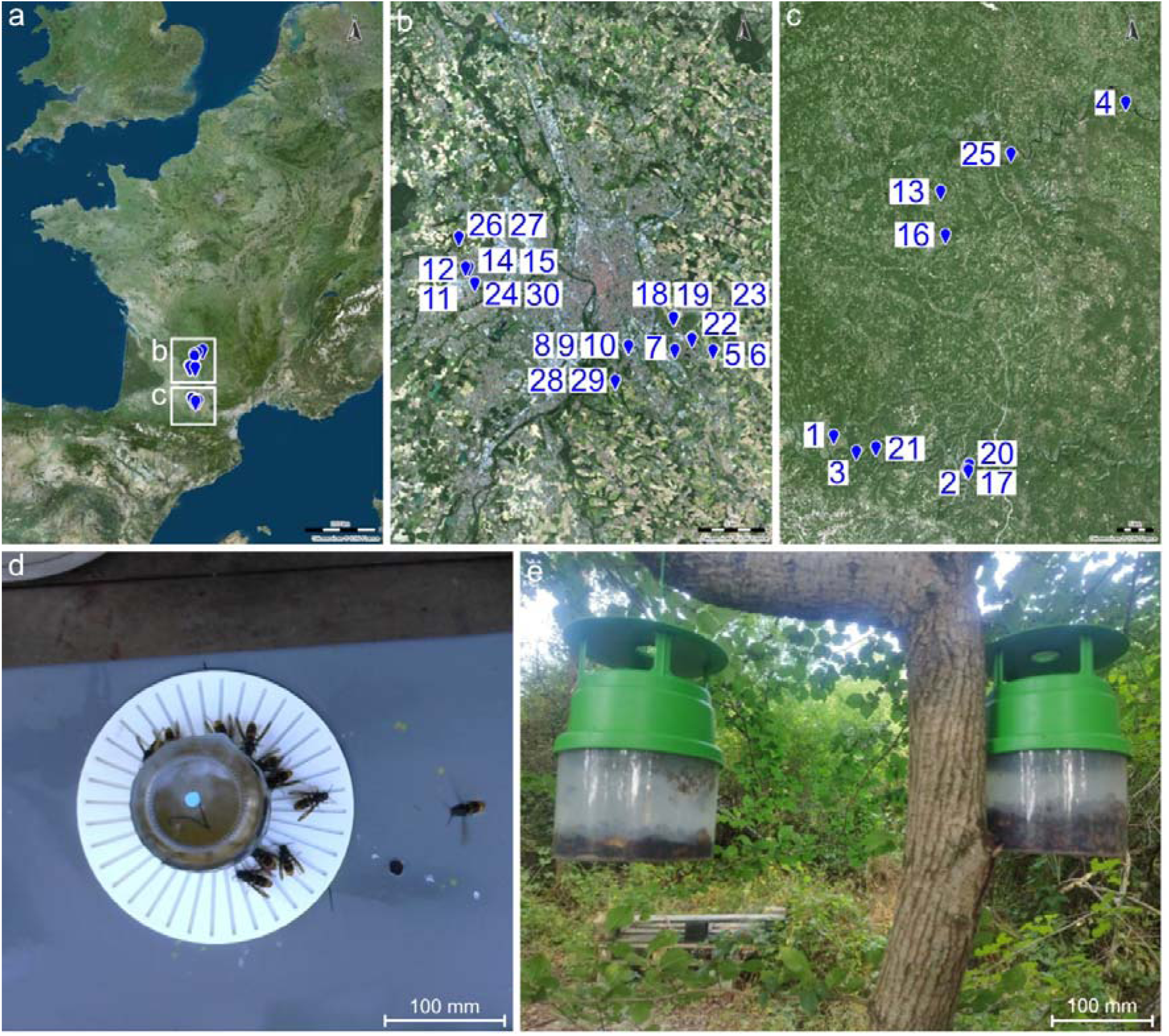
(a) Location of the 30 experimental sites (n°1-30 see details in Table S1; bottom right scale corresponds to 200 kilometers) near (b) Toulouse (scale: 5 kilometers) and (c) Cahors (scale: 5 kilometers). Each site was equipped with two gravity feeders or two baited traps. The feeders (d) were made of a plastic cylinder fitted with a container that released diluted honey solution through gravity. The baited traps (e) consisted of a transparent plastic bowl (diameter: 136 mm; height: 193Cmm; weight: 169 g) covered with an opaque lid (diameter: 16 mm), containing wine, beer and sugar syrup. The traps were placed 50 cm apart, hanging from a tree branch. The maps were created with IGN, Open Licence Etalab 2.0.

## Methods

### Experimental sites

We conducted the study in France over two field seasons in 2022 and 2024. We ran the experiments in 30 sites located in 12 apiaries of comparable sizes (Figure 1, see Tables S1 and S2 for details about site assignment to each experiment). These sites host wild populations of the invasive yellow-legged hornet (*V. velutina nigrithorax,* hereafter referred to as VV). They are also populated by the native European hornet (*Vespa crabro*, hereafter referred to as VC) that occupies a similar ecological niche. Each year we made the observations between June and November in order to exclusively test hornet workers [23].

### Feeder choice experiments

#### Choice between two identical feeders

We tested for inter-attraction in nectar foraging hornets by placing two identical feeders next to each other (hereafter feeder 1 or 2). In case of inter-attraction, we expected one feeder to be chosen by most hornets. We also expected that the frequency of asymmetrical distributions between feeders would increase with the total number of hornets on the two feeders at a given time (group size), and that the position of the feeder attracting the most individuals (left or right, hereafter position A or B) would vary across replicates.

In each site, we positioned two gravity feeders 50 cm apart on a 1m high table (Figure 1d). The feeders consisted in a glass jar (diameter: 5 cm, height: 10 cm) containing liquid food (1:3 ratio of commercial honey (wild flowers honey, Famille Michaud & Co, Gan) to water), placed upside down on top of a 3D printed plastic white landing platform (diameter: 20 cm, height: 0.5 cm) with radial trenches from which hornets could drink. We estimated that each landing platform was large enough to host a group of up to 265 hornets simultaneously. This calculation was made based on the total surface of the landing platform, assuming hornets were elliptical objects 1.98 cm long and 0.57 cm wide [24]. Note that this is an underestimation as hornets could occasionally access food while landing on top of each other. Each jar was covered with dark tape so that the foraging hornets could not see the volume of food available in the feeders upon making a choice. With this design, foraging hornets could choose a feeder based solely on olfactory, tactile and visual cues provided by other hornets already landed on the platforms. During the observations, we swapped the positions of the two feeders every 15 minutes to make sure that attraction to a feeder was solely due to social cues, and not to the identity of the feeders (1 or 2) or to their spatial positions (A or B). The feeders remained at a given location for 30 consecutive minutes (15 minutes in position A and 15 minutes in position B). We ran these experiments in five different locations at one experimental site (site 8 in Table S1) during for four days (Figure 1b), totaling 30 replicates.

#### Data collection on feeders

For each replicate, we recorded the number of hornets on each feeder every 30 seconds using two cameras (C270 HD Webcam, Logitech) placed 60 cm above each feeder. The cameras were operated by a Python3 program [25] using *vidgear* library [26] that automatically took pictures continuously (30 min × 4) between 1 and 6pm (240 frames for 2 hours, 1920 frames in total). We used computer vision with Python’s OpenCV library to quantify the number of darkened pixels (i.e. hornet) on the white surfaces of the feeders’ landing platforms. This involved using adaptive binarization to convert specific luminosity thresholds into black (for hornets) and white (for the background and feeder). To isolate the counting to the feeder area, we applied circle detection tools available in OpenCV. To convert the number of darkened pixels into an estimate of the number of hornets, we used a calibration curve (a polynomial fit of the number of computer-retrieved black pixels throughout a sample of images where the number of hornets was counted by an experimenter (mean error ± sd: 6.99 ± 8.34). We denoted hereafter the total number of hornets in both feeders at a given time as “group size”.

### Trap choice experiments

We tested whether the observations made on gravity feeders could be verified in another context, across more experimental sites and with a larger number of hornets, by giving hornets a choice between two baited traps.

In each experimental site, we placed two commercial hornet traps (Smart Trap®, M2i Biocontrol) 50 cm apart (Figure 1d; Figure S1D). The two traps were hung on the same tree branch, between 1m and 3m above ground. Each trap was made of a transparent bowl containing standard attractive bait (white wine 50% v/v, beer 25% and 25% sugared syrup) and covered with an opaque funnel-shaped lid with a roof. The lid enabled the insects to enter the trap but not to exit. With this design, the hornets could use both olfactory and visual social cues to choose a trap. The traps had low selectivity levels since we caught hornets of two species (i.e. VV and VC) and a variety of other insects including Hymenoptera, Lepidoptera and Diptera (see details in the results).

#### Choice between two identical traps

We first tested for inter-attraction by placing two identical baited traps (traps 1 and 2) next to each other. If the foraging hornets were attracted by trapped hornets, we would expect one of the two traps to be chosen by most individuals. The frequency of significant asymmetries should vary with overall group size (number of hornets in both traps), and the position of the trap attracting the most individuals (A or B) should vary across replicates.

#### Choice between an empty trap and a trap primed with dead hornets of the same species

We then tested whether the social cues involved in inter-attraction were displayed by dead hornets or whether the hornets already present in the traps needed to be alive. Here hornets were given a choice between an empty trap (containing no hornet) and a primed trap (containing 15 dead VV hornets randomly sampled in dedicated traps collected every day and cleaned with fresh water before the experiment). If the dead hornets displayed social cues, we would expect the primed trap to be more frequently chosen by most individuals.

#### Choice between an empty trap and a trap primed with dead hornets of another species

In this final experiment we tested whether the social cues involved in inter-attraction were specific or not. Hornets were given a choice between an empty trap (containing no hornet) and a primed trap (containing 15 dead VC hornets randomly sampled in dedicated traps collected every day and cleaned with fresh water before the experiment). If the inter-attraction by VV hornets was specific, we would expect one of the traps to be chosen randomly by the majority of individuals, irrespective of the presence of dead hornets in the traps.

#### Data collection in traps

We counted the number of hornets in the traps between from 3 to 14 days after setting the traps on site (Tables S1 and S2). We voluntarily varied this duration in order to maximize variance in the total amount of hornets trapped in our dataset, i.e. group size (range: 1 - 422 individuals). For each record, we emptied the two traps simultaneously and counted the total number of VV and VC hornets in each trap. We also estimated the amount of other insects caught in the traps as a percentage of the bowl occupied in order to evaluate their potential influence on hornet attraction.

### Statistical analysis

We performed all the statistical analyses with R 4.5.0 [27]. The raw data are available in a github repository (https://github.com/lacmathilde/Raw_data.git).

For all experiments, at each location of each site, we considered the feeder or trap that attracted the largest number of hornets on each observation period as the “winner” option.

In the choice experiment between two identical feeders, we tested inter-attraction by running a proportional test on each count. We first evaluated if the proportion of hornets in the winner feeder was significantly different from random (i.e., equal proportion) using the function binom.test (*stats* package [27]. We then ran a test to assess whether hornets had accumulated on one of the feeders in a non-random way considering all the counts using the function prop.test. Finally, we assessed the influence of group size on the frequency of asymmetric distributions between two identical feeders by performing a Generalized Linear Mixed-Effects Model (GLMM) with a binomial distribution using glmer function (lme4 package [28]). We used the same statistical analyses for the choice experiment between two identical traps.

In the choice experiment between an empty trap and a primed trap, we tested the influence of dead hornets on captures. We evaluated whether the primed trap was more attractive to hornets and to what extent it more often becomes the winner trap. We then tested if the proportion of captures across the winner and primed traps was different from 0.5 by using a proportional test on each count.

In all trap choice experiments, we evaluated the specificity of social attraction by calculating the proportion of VV and VC hornets caught in the same winner trap. To test whether hornet preys (Vespidae, Apidae, Lepidoptera and Diptera) [21] also caught in the traps influenced the observed distributions of hornets, we performed a linear mixed effect model (LMM) with Poisson distribution using the function lmer (*lme4* package [28]) using *the percentage of trap occupied by the mass of captured insects* as fixed factors and *site* as random factor. We then ran an ANOVA on this model using the function Anova (*car* package [29]).

## Results

### Choice between two identical feeders

We tested the existence of inter-attraction between foraging hornets by giving hornets a choice between two feeders containing *ad libitum* diluted honey. Over the 16 hours of observations (1920 photos), we exclusively observed VV hornets and the group size (total number of hornets landed on the two feeders at a given time) varied between 0 and 215 individuals.

The frequency of asymmetric distributions between the two feeders at a given location increased with group size (Figure 2; GLM with binomial distribution, *Group size* on *asymmetry*: df = 934, z = 5.365, p < 0.001). Accordingly, for small groups of less than 50 hornets, asymmetric distributions were observed in 36.32% of the cases (93 out of 256 measures). This percentage remained stable at 22.48% (29 out of 129 measures) in groups of 50 to 99 hornets, but increased to 67.53% (52 out of 77 measures) in large groups of 100 to 149 hornets, and reached a maximum of 87.50% (7 out of 8) in very large groups of 150 to 199 hornets (Proportional test: X² = 19.97, df = 3, p < 0.001; see pairwise comparisons in Table S3). This group size effect on the frequency of asymmetrical distributions is the signature of a collective decision driven by inter-attraction of insects on feeders.

**Figure 2:**
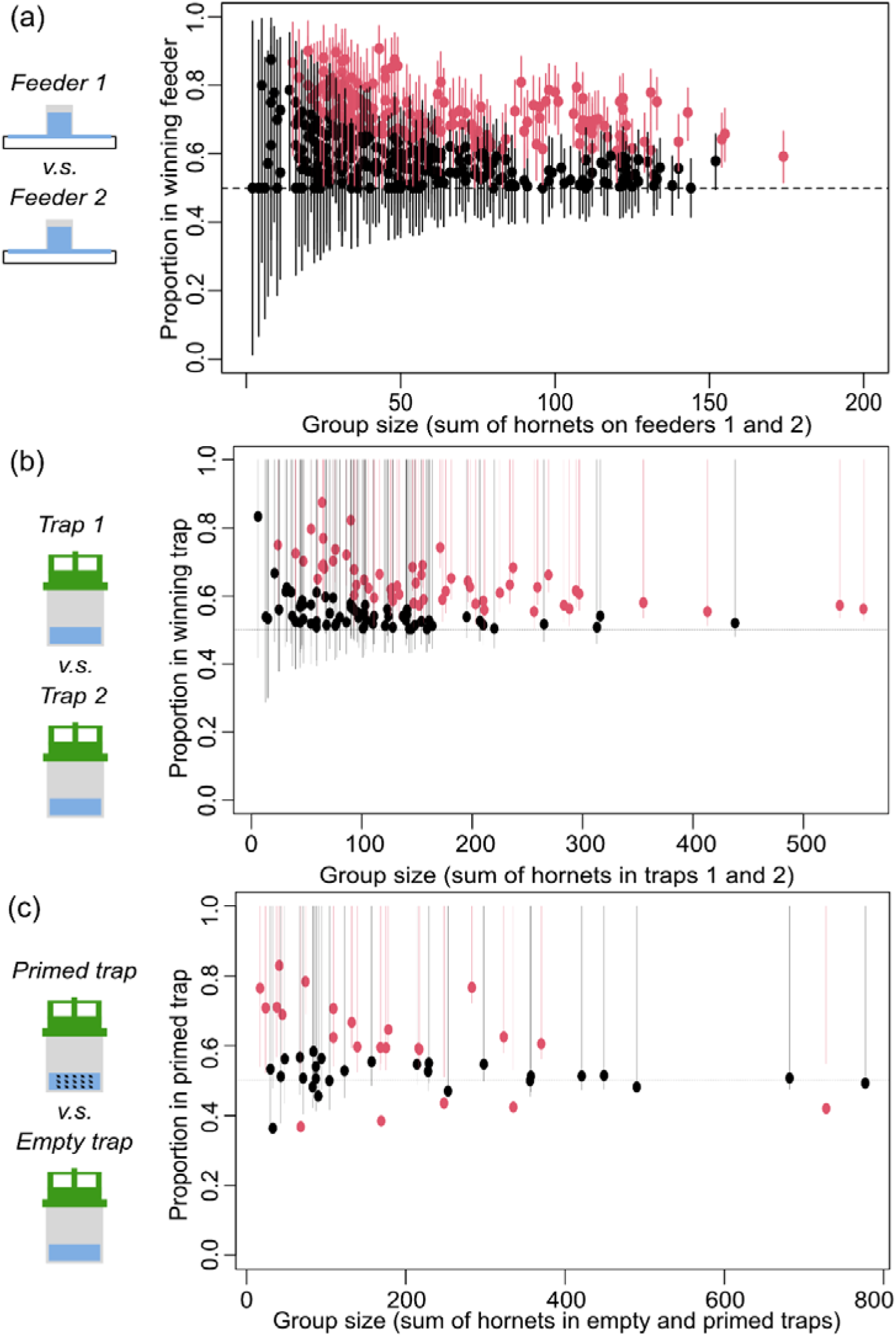
Proportion of hornets in (a) the winner feeder, (b) the winner trap, and (c) the primed trap. Red dots represent statistically significant asymmetric distributions between the two options (proportional test, p<0.05). Black dots are non-significant asymmetries. Solid lines are 95% confidence intervals.

Interestingly, when the distributions of hornets were significantly asymmetric, we observed frequent changes in the identity of the winner feeder through time (Figure 3). Overall, feeder 1 was the winner feeder in only 52.46% of the records (181 out of 345 measures), which means hornets did not have any preference for one of the two feeders (binomial test, p = 0.39). This temporal alternance of winner feeders indicates that inter-attraction was not mediated by long-lasting cues left by hornets on the feeders, such as chemical traces, but rather suggests the implication of visual cues or short-range odours carried by the hornets.

**Figure 3:**
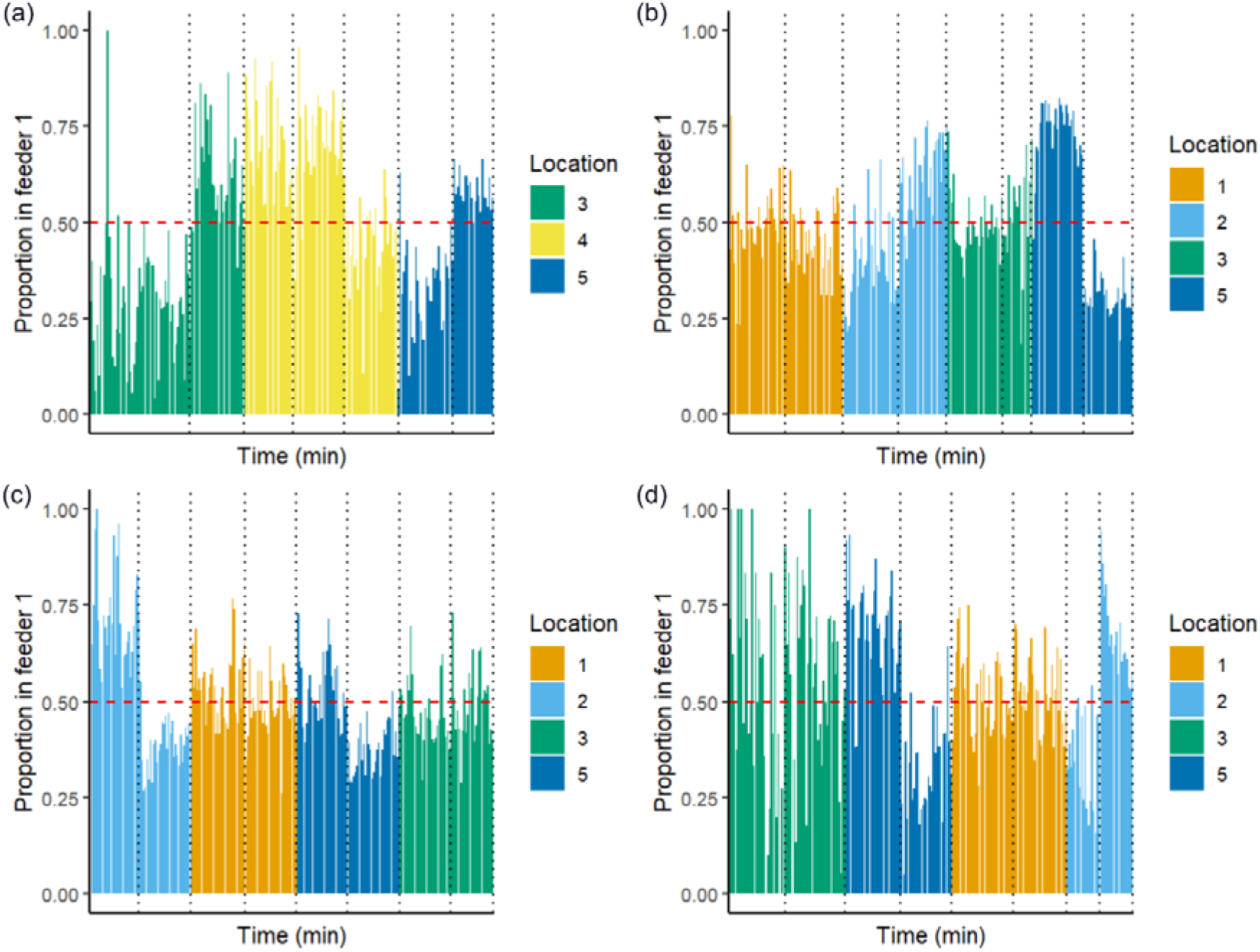
Proportion of hornets on feeder 1 across experimental day (Time) (as: day 1, b: day 2, c: day 3, d: day 4) and in different locations (orange: location 1, light blue: location 2, green: location 3, yellow: location 4, dark blue: location 5) of experimental site 8. The dashed red line represents the point when more than half of the hornet population is located on feeder 1, making it the winner feeder (feeder 1 wins when the proportion of hornets on feeder 1 > 0.5, feeder 2 wins when the proportion of hornets on feeder 1 < 0.5). The dashed black line represents the changes of feeders positions (A or B) every 15 minutes.

### Choice between two identical traps

We tested whether the collective decisions observed for feeders could also be observed for baited traps. This approach enabled us to test more individuals and in more sites.

Here we caught hornets of two species: VV and VC. When considering both species together, the frequency of asymmetrical distributions of hornets among traps increased with the total number of hornets in the two traps (group size) (GLM with binomial distribution, *group size* on *asymmetry*: df = 123, z = 3.12, p < 0.001). The number of records in which the proportion of hornets in the winner trap was significantly higher than in the looser trap increased with group size irrespective of the site, from 31.48% (17 out of 54 records) in small groups with less than 100 individuals in both traps, to 55.56% (40 out of 72 records) in large groups of 100 or more hornets in both traps (proportional test: X² = 6.28, df = 1, p = 0.01). This group size effect was comparable to that previously observed with the feeders (proportional test, group size < 100: X² = 0.002, df = 1, p = 0.96, group size ≥ 100: X² = 1.610, df = 1, p = 0.205).

If inter-attraction were specific, the asymmetrical distributions of VV should be independent of the distribution of VC in the traps. However, when considering the two species separately, we found that in 68.25% of the cases (86 out of 126 records), the winner trap was the same for VV and VC (Figure 2). This was further supported by the fact that traps also caught a diversity of other non-hornet insects, and this number was positively correlated to the total number of hornets of both species in the traps (LMM, percentage of insect mass: X² = 172.1, df = 1, p < 0.001).

### Choice between an empty trap and a trap primed with hornets

To further explore the nature of the attraction cues driving the collective decisions of hornets, we primed one trap with 15 dead hornets and compared the number of hornets caught in the two traps on the same location. If dead hornets displayed attraction cues, we would expect the primed trap to become the winner trap more often than expected by chance.

When considering hornets from the two species altogether, the primed trap was the winner trap in 82.35% of the cases (42 out of 51 records; Figure 2). This confirms that the choice of a trap is influenced by the presence of hornets already caught inside being dead or alive.

If attractive cues were specific, we would expect traps baited with one species to be the winner trap only for that particular species. Accordingly, the VV primed trap became the winner trap for VV in 100% of the cases (39 out of 39 records; binomial test: p<0.001, confidence interval (CI): [0.91; 1.00]). The VV primed trap also became the winner trap for VC overall but in only 67.85% of the cases (19 out of 28 records; binomial test: p=0.09, CI: [0.47;0.84]; 5 records showed equal distribution between the primed and the empty traps). Similarly, the VC primed trap became the winner trap for VC in 100% of the cases (9 out of 9 records; binomial test: p=0.004, CI: [0.66; 1.00]) and the winner trap for VV in only 66.66% of the cases (6 out of 9 records; binomial test: p=0.50, CI: [0.30; 0.93]).

Without distinguishing between caught hornet species, the efficiency of the traps baited with either VV or VC was comparable (χ² test, χ² = 0.001, df = 1, p = 0.924). Indeed, primed traps became winner traps in 84.61% of the cases (33 out of 39 records) when primed with *VV* and 75.00 % of the cases (9 out of 12) when primed with *VC*. Although the inter-attraction appears to be stronger within the same species, both species were attracted to each other.

## Discussion

Collective decision-making is a hallmark of animal sociality [1]. Here, using binary choice experiments in the field, we report collective foraging decisions for food resources in invasive populations of the yellow-legged hornet *V. velutina nigrithorax*. The gregarious behaviours of hornets on feeders and in baited traps is density dependent, and can involve hornets of other species, suggesting that it results from a simple inter-attraction mediated by short range cues passively displayed by foraging hornets.

Our observation of density-dependent asymmetrical distributions of yellow-legged hornets given a choice between identical food sources is a strong signature of collective animal decisions mediated by inter-attraction [8,10,30]. In our experiments, significant asymmetries on feeders containing a sweet solution more than doubled in frequency with group size, from 36.32% when the number of individuals was below 50 to 87.50% when this number reached 150 or more individuals. Critically, a similar density-dependent collective response was observed in hornet populations facing two identical traps also baited with a sweet solution, suggesting that the mechanisms involved in the two aggregation processes are the same. While our results do not enable us to identify potential critical densities above which collective decisions systematically emerge (i.e. quorum), experiments using primed traps show that 15 dead hornets are sufficient to bias the hornet distribution towards the primed trap in ca. 80% of the cases, irrespective of the hornet species used as attractor.

In contrast to the well-documented collective decisions of ants, bees and some other social insects in which individuals attract nestmates on site using active recruitment behaviours [11,12], hornets did not display evident active signaling through specific social interactions or long-lasting scent marking of feeders, as revealed by frequent alternances in the identity of winner feeders during the course of our experiments. Inter-attraction is rather likely mediated by short-range cues passively displayed by hornets on feeders and in traps. Assuming the same mechanism was at play in all our experiments, the fact that hornets in search for food could not interact with trapped insects before entering the actual trap, rules out the implication of tactile cues. The social cues are therefore likely chemical and/or visual. In our experiments, since baits primed with dead yellow-legged hornets successfully biased the collective choices, as well as did dead European hornets to a lesser extent, the attractive cues are not entirely specific and may be used by a variety of species exploiting similar food sources, a phenomenon well-described described in pollinators that can use social information from various species in order to locate most profitable nectar sources [31].

However, at this stage we cannot definitely rule out the possibility that accumulation on feeders and on traps was mediated by different processes and cues, and for instance that hornets were attracted to the traps by cues indicating the presence of potential preys (although we are not aware of any report of consumption of dead hornets by other hornets). Future experiments are therefore needed to further explore the nature of these attractive cues, for instance by manipulating the presence of odours and the visual information available on feeders and in traps.

Irrespective of the precise mechanisms mediating inter-attraction, collective foraging in hornets may provide important benefits for resource localization and exploitation, through information transfer and swarm intelligence, as described in other social animals [32]. First, the fact that yellow-legged hornets were observed to use social information provided by another hornet species may constitute an advantage regarding food localization. For instance, generalist bees, such as bumblebees and honey bees, are well known to use social information from other pollinators to learn about plant species and foraging techniques to obtain the best nectar and pollen resources in their shared habitats [33]. In Western Europe, yellow-legged hornets and European hornets are often seen foraging at in the same areas at similar times of the day [34,35]. They may thus mutually benefit from the presence of each other in terms of information gain regarding plant nectar localization. More generally, collective foraging could enable hornets to adapt to a wide range of habitats by capitalizing on signals already present in their surroundings, ultimately leading to different species benefiting from the presence of each other through heterospecific collective behaviour, as previously documented in other ecological associations [36–39]. Sharing foraging information within and between species may further explain their ecological success and rapid spread in newly colonized areas allowing them for enhanced tracking of most profitable resources in their environment, increased foraging efficiency and higher adaptability to complex environments [36,40,41]. Whether the same collective behavior can be observed in hornets hunting for preys is an open question that will deserve further investigation.

This new observation about the foraging ecology of a major invasive species worldwide provides interesting insights into the context of population control. In particular, this raises the possibility of exploiting the inter-attraction underpinning collective hornet decisions to increase the efficiency of traps by baiting existing trap designs with dead hornets or lures displaying key attractive cues. While many beekeepers already implement this practice based on empirical observations. Our data provide evidence for this phenomenon and offer quantitative predictions related to the effect of group size.

## Acknowledgements

This work was funded by a grant from the French Agency for Ecological Transition (ADEME project LOTAPIS) and the European Research Council (ERC Cog BEE-MOVE, grant number 101002644). We thank Olivier Fernandez for lending its apiaries. We also thank Hugo Cormier, Juliane Mailly and Kristine Abenis for their help in data collection.

## Contribution statement

MLa, DT, FV and MLi designed the study. MLa, BCM, EF, AB, FV, HC, LG collected the data. MLa, CD and JG analysed the data. MLa wrote the first draft. All authors contributed to revisions.

