## Supplementary materials for "Social attraction mediates collective foraging decisions in invasive hornets"

Table S1: Summary table of each counting information to identify an inter-attraction between hornets (two identical traps)

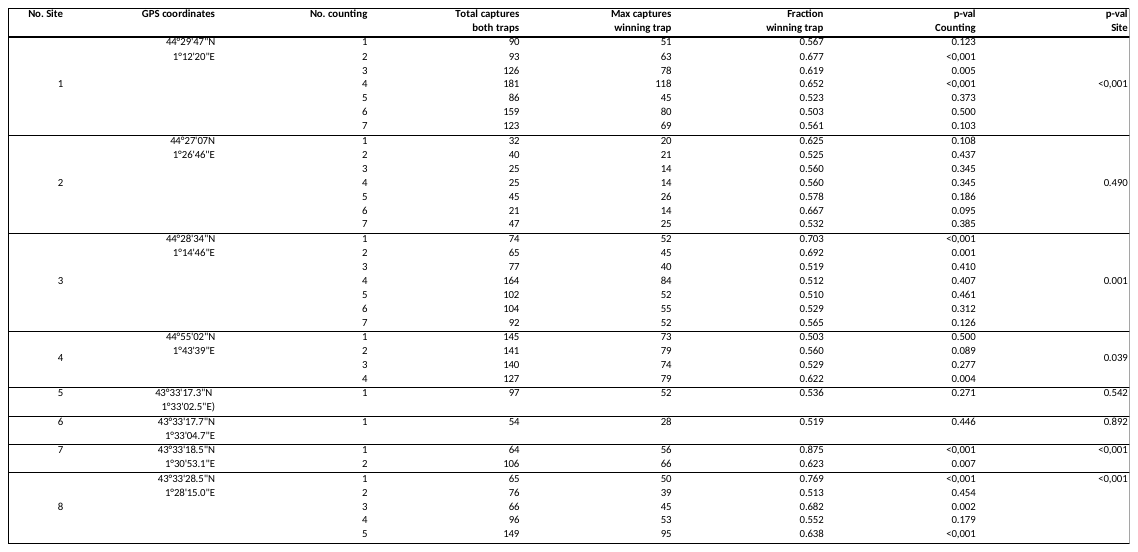

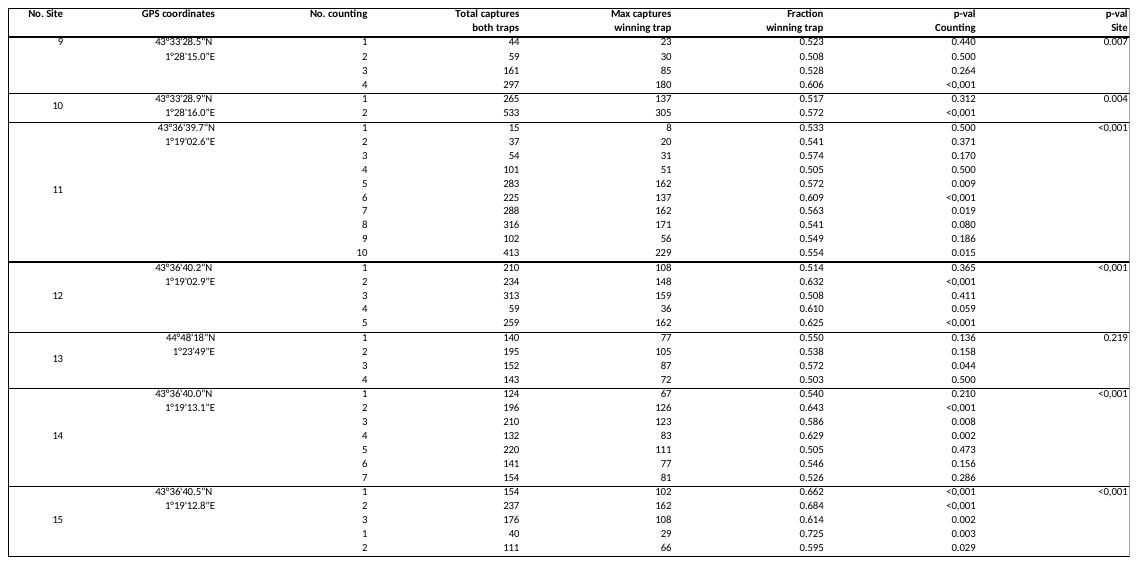

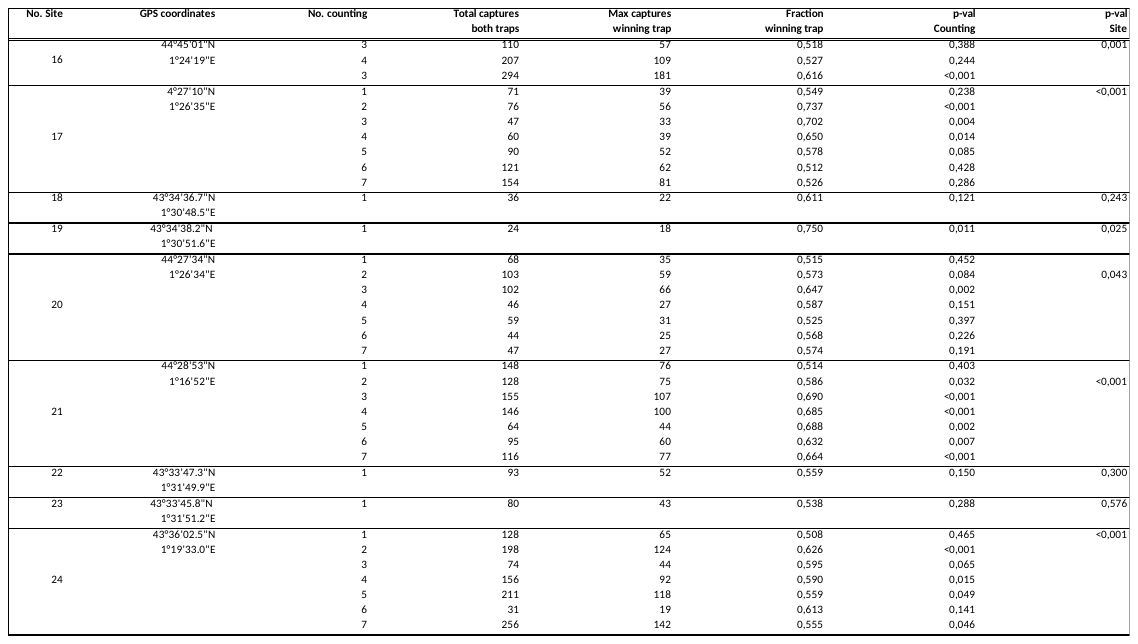

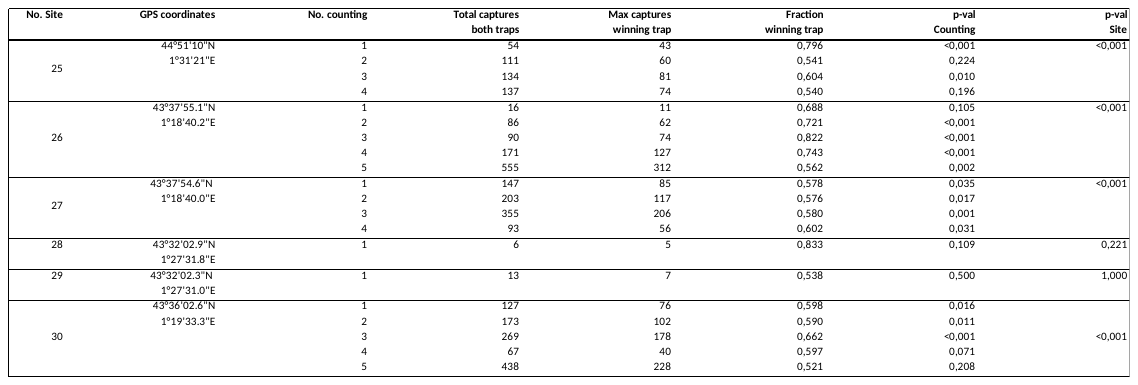

Table S2: Summary table of each counting information to identify the specificity of the inter-attraction between hornets (one primed trap and one emptied trap)

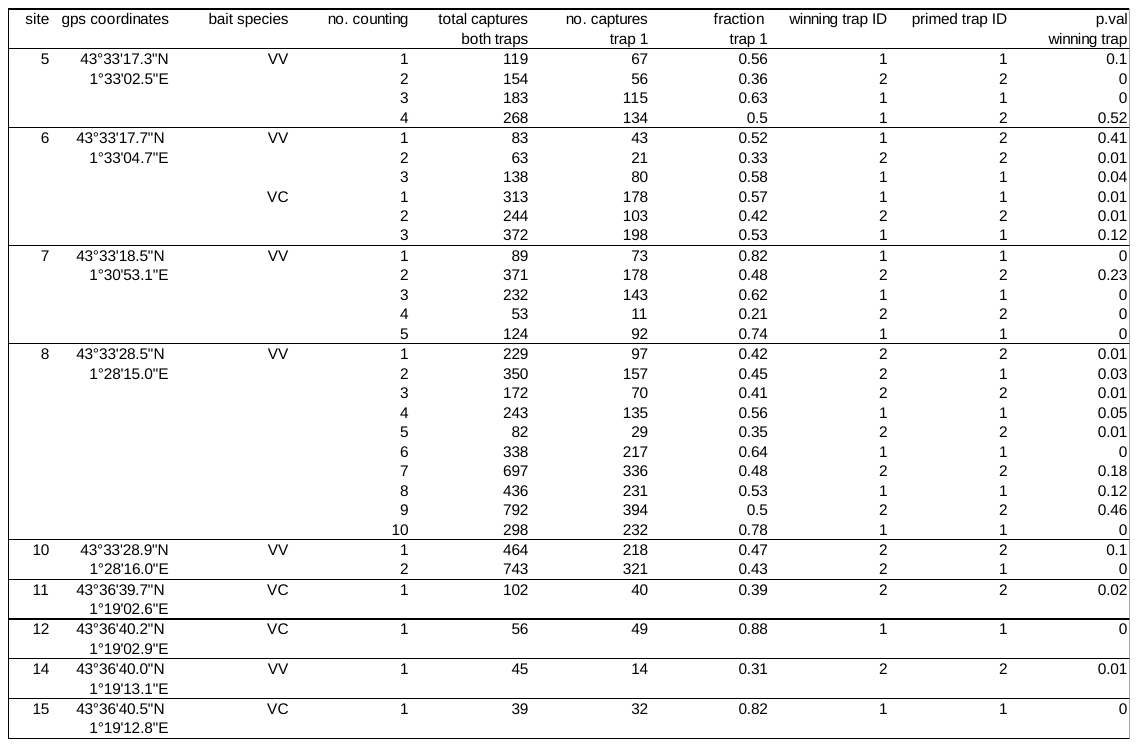

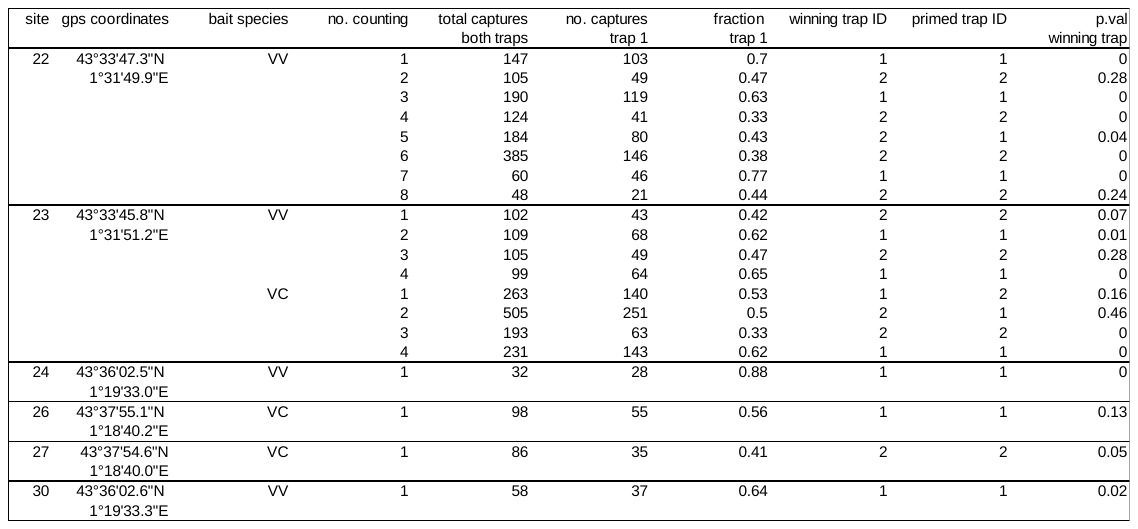

Table S3: Statistical results from pairwise comparison tests with a Bonferroni correction between the proportion of significant asymmetry of different hornets’ group size (designated as GS XX-XX hornets).

|  | **GS 0-49** | **GS 50-99** | **GS 100-149** |
| --- | --- | --- | --- |
| **GS 50-99** | X²= 5.40, df=1,p=0.13 | - | - |
| **GS 100-149** | X²= 23.2, df=1, p<0.001 | X²= 36.80, df=1, p<0.001 | - |
| **GS 150-199** | X²= 5.13, df=1, p=0.14 | X²=9.90, df=1,p=0.001 | X²=0.27, df=1, p=1.00 |
